# Dopamine 2 receptor signaling controls the daily burst in phagocytic activity in the mouse retinal pigment epithelium

**DOI:** 10.1101/789917

**Authors:** Varunika Goyal, Christopher DeVera, Virginie Laurent, Jana Sellers, Micah A. Chrenek, David Hicks, Kenkichi Baba, P. Michael Iuvone, Gianluca Tosini

**Affiliations:** Neuroscience Institute and Department of Pharmacology and Toxicology, Morehouse School of Medicine, Atlanta, Georgia, USA; Institut des Neurosciences Cellulaires et Intégratives (INCI), CNRS UPR3212, 5 rue Blaise Pascal, 67084 Strasbourg, France; Department of Ophthalmology and Eye Center, Emory University, Atlanta, Georgia, USA; Department of Pharmacology, Emory University, Atlanta, Georgia, USA

**Keywords:** Dopamine, Dopamine receptors, RPE, Retina, Circadian, Phagocytosis

## Abstract

**Purpose:** A burst in phagocytosis of spent photoreceptor outer fragments by retinal pigment epithelium (RPE) is a rhythmic process occurring 1-2 hours after the onset of light. This phenomenon is considered crucial for the health of the photoreceptors and RPE. We have recently reported that dopamine, via dopamine 2 receptor (D_2_R), shifts the circadian rhythm in the RPE.

**Methods:** Here, we first investigated the impact of the removal of D_2_R on the daily peak of phagocytosis by RPE and then we analyzed the function and morphology of retina and RPE in the absence of D_2_R.

**Results:** D_2_R KO mice do not show a daily burst of phagocytic activity after the onset of light. Also, in contrast to control, phosphorylation of FAK did not increase significantly in KO mice at ZT1. RNA sequencing revealed a total of 394 differentially expressed genes (DEGs) between ZT23 and ZT1 in the control mice, whereas in D_2_R KO mice, we detected 1054 DEGs. Pathway analysis of the gene expression data implicated integrin signaling to be one of the upregulated pathways in control but not in D_2_R KO mice. No difference in retinal thickness, visual function, or morphology of RPE cells was observed between WT and D_2_R KO mice at the age of 3 and 12 months.

**Conclusions:** Our data suggest that removal of D_2_R prevents the burst in phagocytosis and a related increase in the phosphorylation of FAK after light onset. The pathway analysis points towards a putative role of D_2_R in controlling integrin signaling, which is known to play an important role in the control of the daily burst of phagocytosis by the RPE. Our data also indicate that the absence in the burst of phagocytic activity in the early morning does not produce any apparent deleterious effect on the retina or RPE up to one year of age.

## Introduction

The retinal pigmented epithelium (RPE) is a monolayer of cells that performs many vital functions for maintaining the health and physiology of retina. The RPE is involved in the daily phagocytosis of shed photoreceptor outer segments (POS) (1). Previous studies have shown that the process of RPE phagocytosis starts with shed photoreceptor outer segment interacting with integrin ligand MFG-E8 (2). The next steps comprise of engagement with ανβ5 integrin receptors (ανβ5; 3,4), activation of tyrosine kinases (ανβ5-associated focal adhesion kinase [FAK]); cell surface receptor MerTK (5,6) and re-localization of Rho family GTPase ‘Rac1’ to recruit F-actin to phagocytic cups (7). A recent genomic study in the mouse has also implicated phosphoinositide signaling in the regulation of the peak of phagocytic activity by the RPE (8). Lack of phagocytosis by RPE leads to accumulation of POS debris in the subretinal space followed by photoreceptors degeneration in the rat (9) and mice (10).

Further studies have also shown that the phagocytic activity of the RPE peaks in the morning, shortly after the onset of light (1-2 hrs) in many different mammalian species (11,12,13,14). This peak or burst in phagocytosis follows a circadian rhythm as the peak continues to occur even after the animals are kept in complete darkness (11,12,13,14). Disruption in the daily peak of RPE phagocytosis impairs retinal and/or RPE functions since mice in which the ανβ5 integrin receptor has been disabled by removing the gene *Itgb5* which codes for the β5 subunit of the receptor (β5^−/−^ mice), fail to show the morning burst of phagocytic activity. Moreover, during the aging process, β5^−/−^ mice also show a decrease in visual function (i.e., the amplitude of the a- and b-wave of the scotopic ERG) and accumulate more lipofuscin in the RPE with respect to control mice (4). Interestingly, it has also been reported that just a change in the timing of the peak in the phagocytic activity is associated with an increase in lipofuscin accumulation and photoreceptor loss during aging (15). Finally, it was worth noting that the mechanism controlling this peak seems to be located within the RPE since the diurnal rhythm, in the exposure of phosphatidylserine by rod outer segments is not entirely controlled by the photoreceptors, but RPE cells participate in the synchronization of this process (16).

Several lines of evidence suggest that dopamine (DA) is involved in the regulation of rhythmic RPE functions in vivo. For example, inhibition of DA synthesis during the early part of the light phase induced a significant reduction of disk shedding and phagocytosis (17) and mice whose dopaminergic neurons have been destroyed by 1-methyl-4-phenyl-1,2,3,6-tetrahydropyridine (MPTP) accumulate a large number of residual bodies in the RPE (18). Besides, we have recently reported that a circadian clock is present in the RPE (19) and this clock is entrained to the external light-dark cycles by DA via dopamine 2 receptors (D_2_R) that are present on the RPE (20).

In this study, we first investigated the effect of D_2_R removal on the daily burst of phagocytic activity and then we explored the effects that removal of these receptors may have on retinal and RPE morphology and function during aging.

## Material and Methods

### Animals

D_2_R (Drd2^tm1Low^) Knock- Out (KO) and C57BL/6J (WT) mice, purchased from The Jackson Laboratory (Bar Harbor, Maine) and bred at Morehouse School of Medicine, were used in this study. Mice were maintained in 12 h light and 12 h dark with lights on (zeitgeber time [ZT] 0) at 06:00 a.m. and lights off [ZT 12] at 6:00 p.m. Genotype of the mice was confirmed by PCR analysis of DNA samples prepared from tail tips by using the primers 5’-CACTCCGCCACTTGACATACA-3’ and 5’-TCTCCTCCGACACCTACCCCGA-3’. All experimental procedures were conducted in accordance with the NIH Guide on Care and Use of Laboratory Animals and were approved by the Morehouse School of Medicine Animal Care and Use Committee.

### Phagosome counting

Eyecups were fixed overnight in 4% paraformaldehyde at 4°C, transferred to series of sucrose solutions (10%, 20%, and 30%, each for 2 hours) and embedded (Tissue-Tek; Sakura Fintek, Tokyo, Japan # 4583). Then 10-*μ*m-thick cryostat sections were prepared and stored at −20°C until ready for use. The sections were permeabilized with Triton X-100 (0.1% in PBS for 5 minutes) and then saturated with PBS containing 0.1% BSA, 0.1% Tween-20, and 0.1% sodium azide (buffer A) for 30 minutes. Sections were incubated overnight at 4°C with primary antibody diluted in buffer A. The primary antibody used for this purpose was monoclonal anti-rhodopsin antibody Rhodopsin 4D2 (see [15] for details). Secondary antibody incubation was performed at room temperature for 2 hours with Alexa (488 nm) goat anti-rabbit (Thromofischer Scientific, Santa Clara, CA # R37116). Cell nuclei were stained with 4,6-diamino-phenyl-indolamine (DAPI; Invitrogen Carlsbad, CA # D1306). Slides were washed thoroughly, mounted in PBS and glycerol (1:1), and observed with a confocal laser scanning microscope (LSM 700 ver. 2.5; Carl Zeiss Meditec, Jena, Germany) (15).

### Western blot

RPE samples were obtained from the eyes of control and KO mice at ZT23, ZT1 and ZT3 using the methodology described in Baba et al., (2010) and then lysed in ice-cold RIPA buffer (50mM Tris, pH 8.0; 150mM NaCl; 1mM EDTA; 1mM EGTA; 1.0 % Nonidet −40; and 1.0 % sodium deoxycholate), 1x protease inhibitors, and 1x phosphatase inhibitor I and II. Following lysis and separation on SDS/PAGE gel, the proteins were transferred to PVDF membrane (Trans-Blot Turbo transfer system; Biorad Laboratories, Hercules, CA #1704156) The blot was incubated overnight at 4°C with Phospho-FAK (Tyr 397) (1:1000, Cell Signaling, Danvers, MA # 3283), FAK (Cell signaling, Danvers, MA, # 3285). RPE-65 (1:2500, generous gift of Dr. T.M. Redmond, NEI). RPE-65 was used as a loading control for the amount of RPE protein present in the samples. Then, they were incubated with anti-rabbit HRP (1: 10000, Cell signaling, Danvers MA # 7074S) and developed. Band intensities were quantified by densitometry using Image J (1.51w).

### Quantitative real-time-polymerase chain reaction (Q-PCR)

RPE samples were collected at ZT23 and ZT1 and subjected to RNA extraction using Trizol (Thermo Fisher Scientific, Santa Clara, CA # 15596026). Q-PCR was performed using the CFX96 Touch Real-Time PCR Detection System (Bio-Rad Laboratories, Hercules, CA, USA) using iQ SYBR Green Supermix (Bio-Rad Laboratories Hercules, CA# 1708880). All data for individual genes were normalized to 18S, according to a previously published protocol (21).

### Library preparation for RNA-sequence (RNA-seq)

Eyes were obtained from WT and D_2_R KO mice at ZT23 and ZT1 (n=3 for each time point). The anterior segment, along with the neural retina, was dissected from the posterior segment that contains the RPE, choroid, and sclera. Following homogenization, by sonicator, the isolated RPE cells were processed for RNA isolation with TRIzol (Ambion, 15596018) following the manufacturer’s instructions. The total RNA was used to prepare 12 mRNA libraries following the standard Illumina protocol. Total RNA samples from the RPE were sent to Omega Bioservices (Norcross, GA) for both library preparation and next-generation sequencing.

### RNA seq runs, mapping and estimation of Reads per kilobase per million (RPKM)

The 12 RNA-seq libraries were then sequenced on the Illumina HiSeq2000 platform at the to produce approximately 65 million, 100 nucleotide paired-end reads per sample (Reads 1 and 2). The reads were mapped to the University of California–Santa Cruz (UCSC; Santa Cruz, CA, USA) mouse genome assembly and transcript annotation (mm10). Mapping was performed with Bowtie2 (v 2.1.0) using the default settings. HTSeq-count (PyCharm Community Edition 2016.3.2) was used to generate counts of reads uniquely mapped to annotated genes using the UCSC mouse assembly mm10 gtf file. Further, RPKM (reads per kilobase per million) were calculated manually and only the genes having an RPKM of ≥ 1 were considered for further analysis (23). Fold change was later calculated by using the RPKM values of the same gene at two different time points (ZT1 *vs.* ZT23). Finally, we used *t*-test (paired-end; ≤0.05) and a cutoff of fold change ≥ 1.5 to determine the differentially expressed genes (DEGs).

To evaluate the significance of the identified DEGs, we used the Protein ANalysis THrough Evolutionary Relationships (PANTHER) Classification System and analysis tools to categorize DEGs by protein class, Gene Ontology (GO), Molecular Function, and GO Biological Process and also to determine if any of these classes or GO terms were overrepresented. First, differentially expressed genes with a fold change ≥2, were identified among the DEG’s in each genotype and entered in the PANTHER database in increasing order of the *p*-value. The PANTHER Overrepresentation Test (release 20171205) was used to search the data against the PANTHER database (PANTHER version 13.1 Released 2018-08-09) and the GO database (Release 20171205) to identify GO annotations and pathways overrepresented in our data when compared to a reference mouse genome.

### Spectral Domain Optical Coherence (SD-OCT)

A Micron IV SD-OCT system and a fundus camera were used (Phoenix Research Labs, Pleasanton, CA) to analyze retinal morphology in the WT and D_2_R KO mice at 3 and 12 months of age. Image-guided OCT images were obtained for the left and right eyes after a sharp and
clear image of the fundus (with the optic nerve centered) was obtained. SD-OCT was a circular scan about 100 μm from the optic nerve head. Fifty scans were averaged. The retinal layers (indicated on the figure images) were identified according to published nomenclature. Total retinal thickness and thickness of the individual retinal layers were analyzed by using NIH Image J (1.51w).

### Electroretinography (ERG)

Mice were subjected to dark adaptation for at least 1 hour and anesthetized with ketamine (80 mg/kg) and xylazine (16 mg/kg). The pupils were dilated with 1% atropine and 2.5% phenylephrine (Sigma, St. Louis, MO, USA). Mice were placed on a regulated heating pad set at 37°C with feedback from the rectal temperature probe. The eye was lubricated with the saline solution, and a contact lens type electrode (LKC Technologies model: N1530NNC) was topically applied on the cornea. A needle reference was inserted in another side of the cheek, and the ground needle was inserted into the base of the tail. All preparation of ERG recordings was conducted under dim red light (<3 lux, 15W Kodak safe lamp filter 1A, Eastman Kodak, Rochester, NY, USA).

All electrodes were connected to a Universal DC Amplifier (LKC Technologies model UBA-4200) and the signal was filtered from 0.3 to 500 Hz. Data were recorded and analyzed by EM for Windows (ver. 8.2.1, LKC Technologies). In the dark-adapted (scotopic) ERG protocol, seven series of flash intensity between from 0.03 to 6.28 cd*s/m^2^ were presented to the mouse eye. Flashes were generated by 530-nm green LEDs in a Ganzfeld illuminator (LKC Technologies), and intervals of flashes increased from 0.612 to 30 s as the intensity of the flashes increased. Responses of 3–10 flashes were averaged to generate a waveform for each step of light intensity, and a-wave and b-wave of ERG measurement were analyzed from the trace of waveforms (22)

To measure the light-adapted (photopic) ERG, mice were placed in a Ganzfeld illuminator and cone-associated activity was isolated by saturating rods with 63 cd*s/m^2^ of steady white background light. The four series of consecutive 10 white flashes (79.06 cd*s/m^2^) were introduced at 2.5 min, 5 min, 10 min, and 15 min during the background light exposure. The Background light was left on for 15 min while photopic ERGs records were measured. The traces of the ERG were averaged and stored on a computer for later analysis. The amplitude of the b-wave was measured from the trough of the a-wave to the peak of the b-wave or, if no a-wave was present, from the baseline to the b-wave peak. The spectral composition and irradiance of the light was monitored by a radio-spectrophotometer (USB 2000, Ocean Optics, Dunedin, FL).

### RPE Morphological Analysis

WT and D_2_R KO mice were euthanized using cervical dislocation; the eyes were collected and then placed in Z-fix (Anatech, #170) for 10 minutes at room temperature and then were rapidly rinsed with 0.1 M phosphate buffer saline five times. Then RPE flat mounts were prepared as previously described (24). Briefly, each eye was placed on a clean microscope slide (VWR, Suwanee, GA # 16004-406) and any excess muscle or fat surrounding the eye was removed with spring scissors (WPI, Worcester, MA # 501235). A single puncture was made in the middle of the cornea with Dumont #5 fine forceps (FST, Foster, CA # 11254-20). By using Dumont #5/45 (FST, Foster, CA # 11251-35) to support the eye globe, spring scissors were inserted into the initial puncture in the cornea and 4 radial cuts were made towards the dorsal end of the eye to where the optic nerve was located. The eyecup was then open and the lens was removed from the center of the eye. Next, the iris and the attached cornea was removed from the anterior ends of the petals. The retina was then removed to expose the underlying RPE. If necessary, center cuts in a petal were made to reduce the overall tensile stress of the RPE sheet.

Flat mounts were then incubated in a 1% bovine serum albumin (BSA, Tocris Biosciences, Minneapolis, MN # 5217) blocking solution for 30 minutes at room temperature. The blocking solution was removed, and the flat mounts were incubated overnight in ZO-1 (Zonula occludens-1) antibody (1:500; Millipore Sigma, Burlington, MA # MABT11) solution diluted in 1% BSA blocking solution at 4 °C. On the next day, flat mounts were washed in a 1x Hank’s Buffered Salt Solution (Gibco, Gaithersburg, MD # 14065-056) with 0.1% Tween-20 (BioRad Laboratories, Hercules, CA # 170-6531) for 5 times at 2 minutes each. Then the flat mounts were incubated in secondary antibody conjugated with Alexa Fluor 488 (1:500; goat anti-rat, Invitrogen, Carlsbad, CA # A11006) for 3 hr at room temperature on a shaker. Flat mounts were washed in the previously mentioned wash solution for 5 times at 2 minutes each. Under a dissecting microscope, the flat mounts were flattened onto slides and mounted with Vectashield antifade mounting medium (Vector Laboratories, Burlingame, CA # H-1000) and allowed to dry at room temperature overnight. ZO-1 signal in the flat mount was visualized using a confocal microscope (Zeiss LSM 700) with two to three images of the peripheral region and four to six images from the central region captured from each petal of the flat mount. Images were then processed in CellProfiler (v2.2.0) to determine the RPE morphological features such as area, eccentricity, compactness, and solidity (24).

### Statistical Analysis

In all cases, the data were tested for normal distribution using the Shapiro-Wilk test and equal variance was tested using the Brown-Forsythe test to check the fulfillment of the requirements for the parametric tests. Data were analyzed with *t*-test, one- or two-way ANOVA. *Post hoc* multiple comparisons of interactions were performed with Tukey’s test. Significance level was set at *p* < 0.05. Data are expressed as the mean ± standard error of the mean.

## Results

### Lack of D_2_R signaling abolishes the daily burst in phagocytic activity by the RPE

Quantification the phagosomes in the retinal sections at ZT23, ZT1 and ZT3 of WT and D_2_R KO mice revealed that phagocytic activity was significantly different between the two groups (n=4; two-way ANOVA; *p* <0.001 for genotype x time interaction). As expected in WT mice the number of phagosomes in the RPE significantly increased at ZT1 and then declined to basal level at ZT3 (two-way ANOVA, followed by Tukey test, *p* < 0.05) whereas in D_2_R KOs RPE phagocytic activity did not show any differences among the three time points (Two-way ANOVA, *p* >0.05; Fig.1). Comparison between both genotypes at all the three time points (ZT23, ZT1 and ZT3) showed that D_2_R KO RPE had significantly fewer phagosomes at ZT1 as compared to WT (two-way ANOVA, *p* < 0.001; Fig. 1).

**Fig 1.**
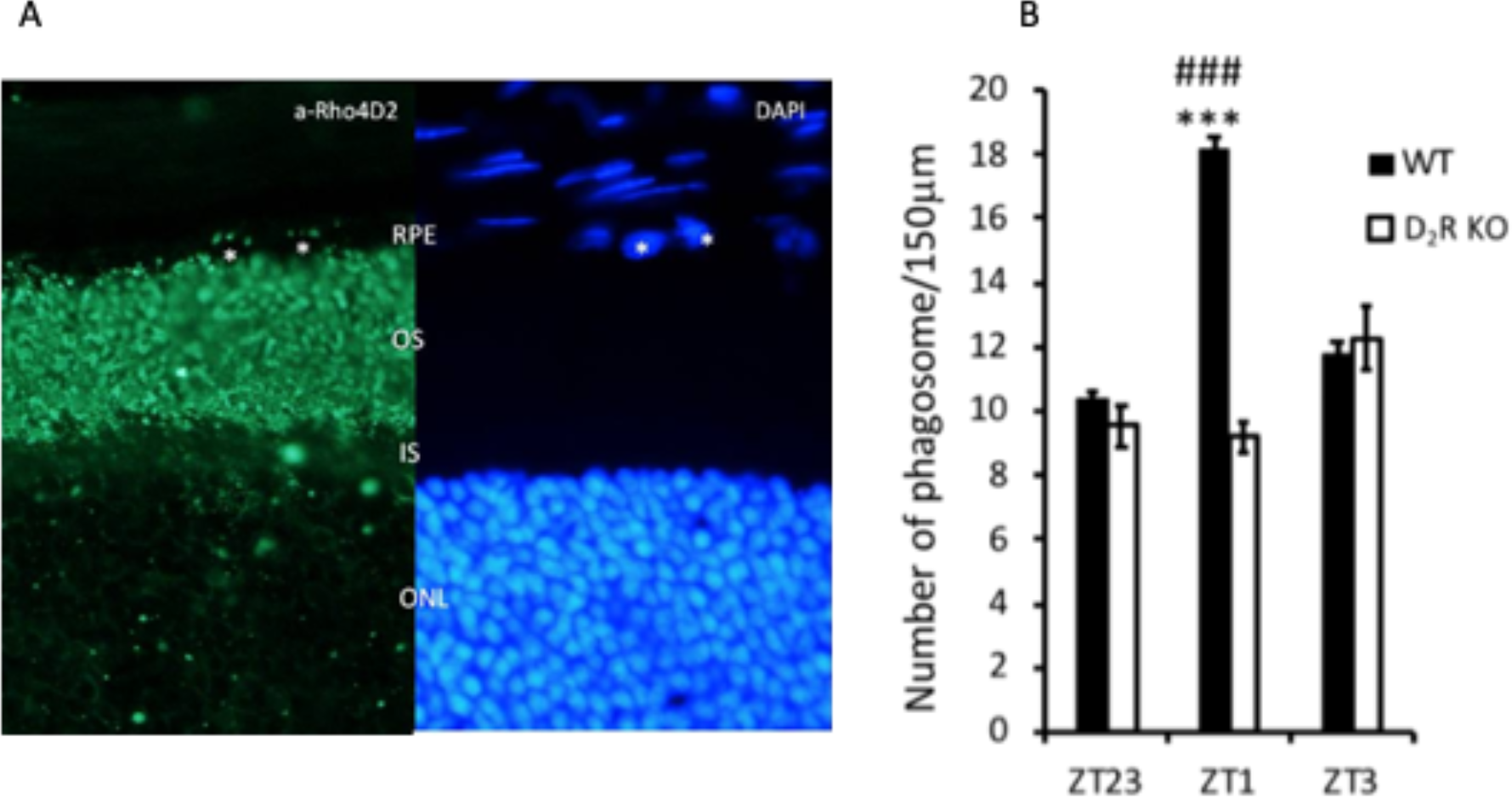
D_2_R KO mice lack the burst of phagocytic activity after light onset. (A) Representative microphotograph of the mouse retinal section immunostained with anti-rhodopsin antibody (Rho4D2, green) and DAPI (blue). Phagosomes (marked with a circle) can be seen as small particles present in RPE co-stained for rhodopsin and DAPI. ONL: outer nuclear layer, OS: outer segment, RPE: retinal pigment epithelium. (B) Bar graph indicating the number of phagosomes/150 /m of RPE at different time points. WT mice showed a significant increase in phagocytic activity at ZT1 (two-way ANOVA, followed by post hoc Tukey test; \*\*\**p* <0.001) while no differences in the number of phagosomes was observed in D_2_R KO at the three different time-points (two-way ANOVA, *p* > 0.1). Bars represent means ± SEM. n=4-6

### Removal of D_2_R inhibits the activation of FAK after the onset of light

Western blot analysis indicated that the level of FAK phosphorylation at Tyr397 (p-FAK) in WT mice was increased at ZT1 with respect to the values observed at ZT23 and ZT3 whereas no change in p-FAK levels was observed in D_2_R KO mice at ZT23, ZT1 and ZT3 (Fig. 2A). Densitometry analysis of the western blot data confirmed that p-FAK level was significantly less at ZT1 in D_2_R KO. Also, no other difference was detected between the two genotypes at the other time points (Fig. 2B, n=3, two-way ANOVA followed by Tukey test). Total FAK levels remained constant and were the same in both the genotypes (two-way ANOVA, *p* > 0.05).

**Fig 2.**
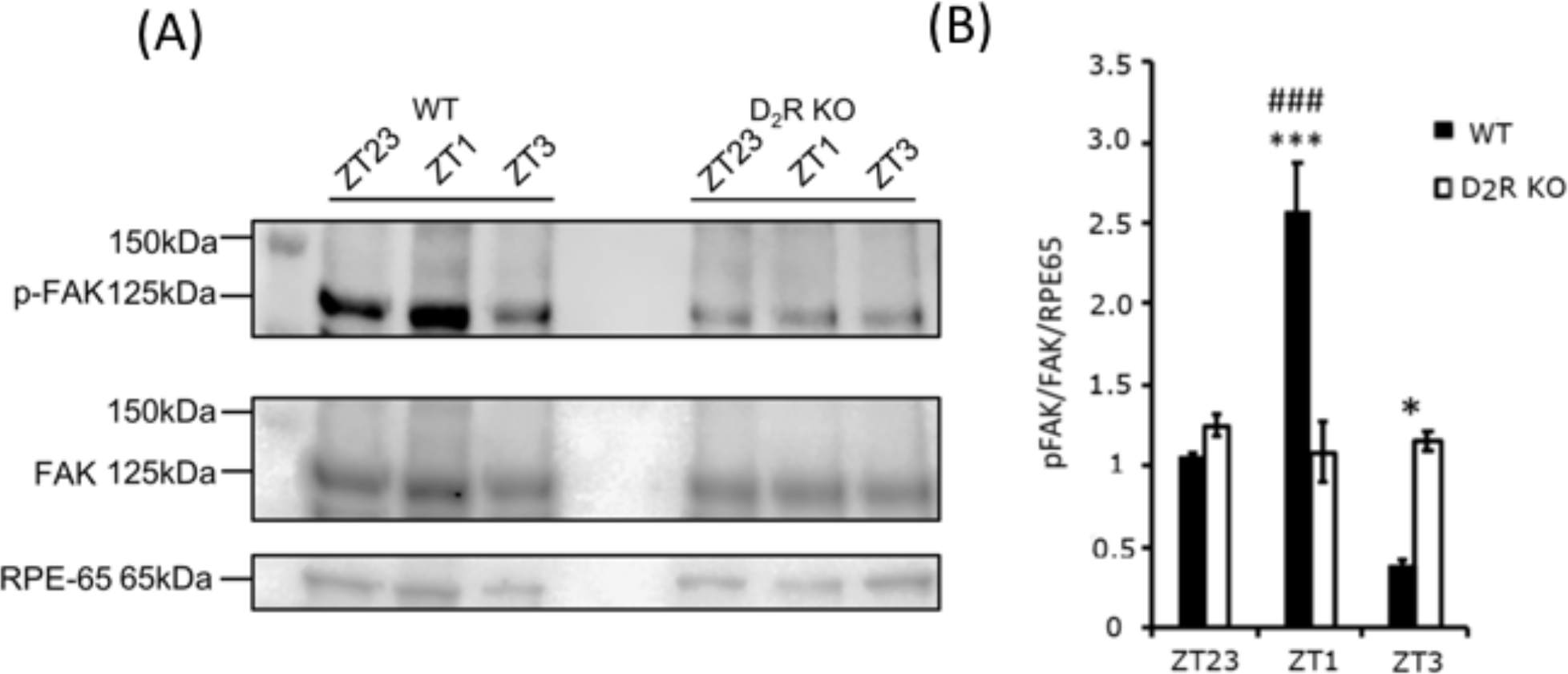
D_2_R KO mice lack p-FAK increase after light onset. (A) Representative western blot bands for RPE-65, FAK and p-FAK in WT and D_2_R KO mice at ZT 23, 1, and 3. (B) Densitometry analysis of the bands intensity indicated a significant increase in p-FAK after the onset of light in WT (two-way ANOVA; \*\*\**p* <0.0001), but not in D_2_R KOs (two-way ANOVA; *p* >0.05). Bars represent the means ± SEM. \**p* <0.001. n=3

### Removal of D_2_R signaling does not affect the expression of clock genes in the RPE *in vivo*

Expression levels of mRNA in the RPE over the course of the day for three canonical clock genes (*Per1*, *Bmal1* and *Rev-Erbα*) was analyzed by quantitative PCR (Fig 3). In both genotypes, the three clock genes showed a rhythmic pattern and no differences in the expression levels or phase were observed between WT and D_2_R KO mice (n= 5-6; two-way ANOVA, *p* > 0.05 in all cases).

**Fig 3.**
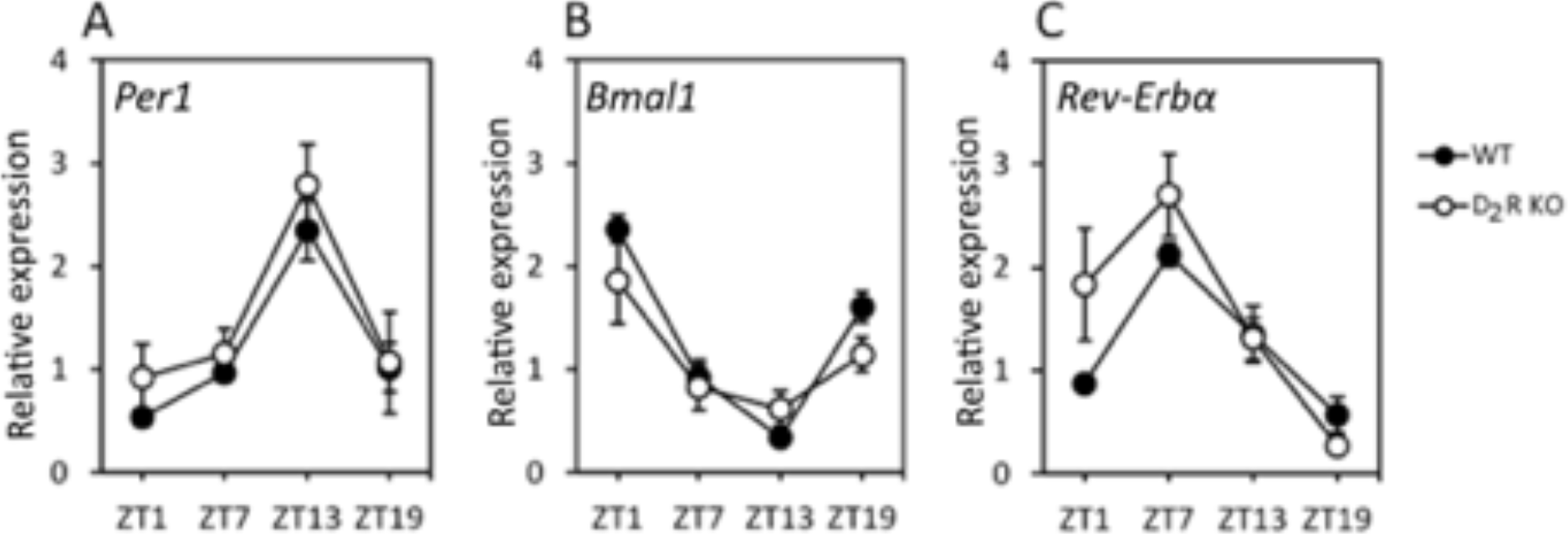
Removal of D_2_R signaling does not affect the expression of the clock genes. *Per1* (A), *Bmal1* (B) and *Rev-ErbR* (C) mRNA expression levels, rhythmic pattern and phase were not affected by the removal of the D_2_R (two-way ANOVA, *p* > 0.1 in all cases). Circles represent the means ± SEM. \**p* <0.001. n=4-6.

### RNA-Seq analysis of the RPE reveals differentially expressed genes between time points in WT and D_2_R KO mice

To identify differentially expressed genes (DEGs) between the different time points (ZT23 *vs*. ZT1) and genotypes, we used a cutoff expression of 1 RPKM, equivalent to one transcript per cell. A total of 13,387 and 12,487 protein-coding gene transcripts were detected in the RPE of WT and D_2_R KO RPE, respectively, between the two time points (ZT23 and ZT1; Table 1). We also observed a higher number of DEGs in D_2_R KO mice (1054 genes) than in WT (394 genes) between ZT1 and ZT23 (*p* ≤0.05). Interestingly only a small number (39) of DEGs overlapped between the two genotypes and although most (36) of the genes showed a common pattern, 3 genes were upregulated in one genotype and down-regulated in the other (Table 1). Full details of the differential expression analysis can be found in supplementary Table 1S and Table 2S and the biological process in which DEGs are involved at the two different time points and in the two genotypes are listed in Fig. 4.

**Table 1.** The table summarizes the gene expression profile in the RPE of WT and D_2_R KO mice between ZT23 and ZT1 time point. Although the total number of transcripts were not different between two genotypes, the number of Differentially Expressed Genes (DEGs) were higher in the RPE of the D_2_R KO mouse than in WT.

|  | WT | D<sub>2</sub>R KO |
| --- | --- | --- |
| Total Transcripts (> 1 RPKM) | 13387 | 12478 |
| Differentially expressed transcripts | 394 | 1054 |
| Upregulated | 247 | 651 |
| Downregulated | 147 | 403 |
| Fold Change > 2 | 183 | 854 |
| Fold change $\geq 1.5 = < 2$ | 211 | 200 |

**Fig 4.**
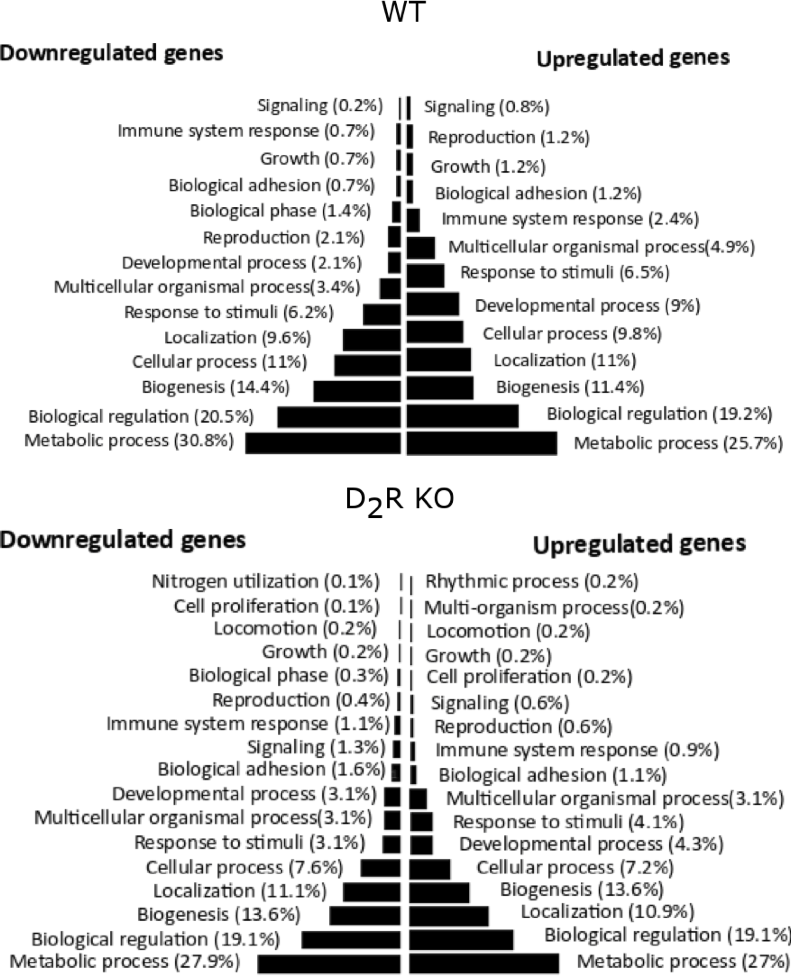
Gene ontology analysis and gene set enrichment in WT and D_2_R KO mouse RPE at ZT23 *vs*. ZT1. The figure shows the biological processes that are up- and down-regulated after the onset of light in WT and D_2_R KOs. In both genotypes, the onset of light-induced an up-regulation of metabolic process, biological regulation and biogenesis.

### Integrin pathway is dysregulated in D_2_R KO mice

Gene ontology analysis indicated the involvement of multiple signaling pathways that were up- or down-regulated in the two genotypes (Fig. 5A, B). In WT mice, the integrin signaling pathway was up-regulated at ZT1, whereas in D_2_R KO mice, this signaling pathway was down-regulated. Further analysis of the genes associated with the integrin signaling pathway revealed significant differences in expression of these genes. In WT mice Rap guanine exchange factor 1 (*Rapgef1*), Jun Proto-Oncogene (*Jun*), Myosin light chain, Phosphorylatable, Fast skeletal muscle (*Mylpf*), and Fyn Proto-Oncogene (*Fyn*) and phosphatidylinositol-4,5-bisphosphate 3-kinase (*Pi3K)* were up-regulated at ZT1 whereas most of these genes were downregulated in D_2_R KO (Fig. 5C, D). Genes like *Itgb5* and *Gsk3b* that were not differentially expressed in WT were down-regulated (more than 2-fold) in D_2_R KO mice (Fig. 5C, D).

**Fig 5.**
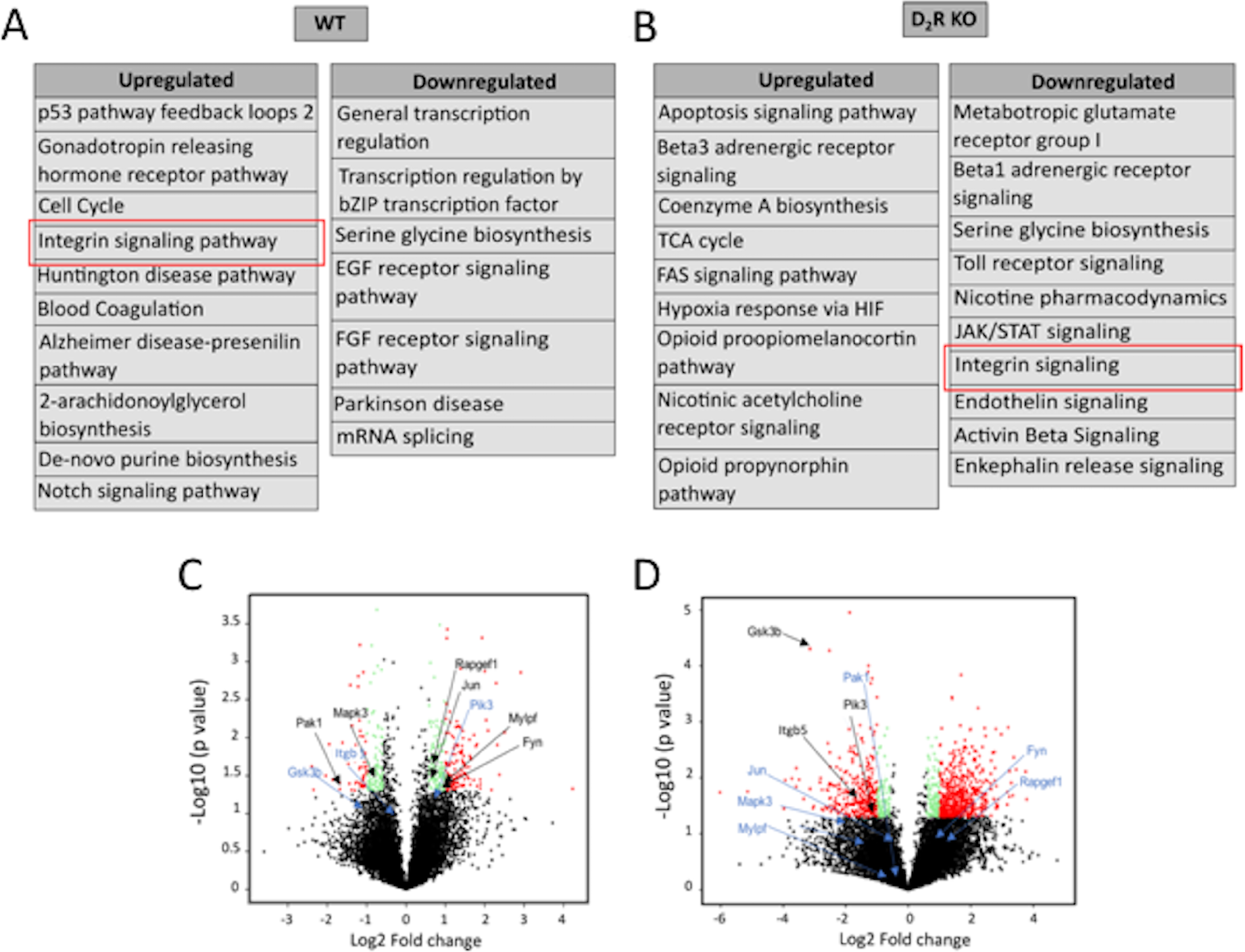
Pathway analysis of differentially expressed genes from WT and D_2_R KO mouse RPE. (A, B) Differentially expressed genes having a fold change of > 2 were analyzed in PANTHER to identify upregulated and downregulated pathways in both genotypes. While integrin signaling (marked in the red box) was upregulated in WT, it was found to be downregulated in D_2_R KO mouse RPE. (C, D) Volcano plot of transcript oscillations between ZT23 and ZT1 in WT and D_2_R KO mouse RPE. The black dots indicate the non-differentially expressed genes (fold change < 1.5 and *p*-value >0.05). Green and red dots indicate the differentially expressed genes having a fold change > than 1.5 and 2, respectively, and a *p*-value < 0.05. Also marked are the genes involved in integrin signaling; upregulated (black) and downregulated (blue).

### D_2_R KO mice do not show any changes in retinal morphology or ERG responses

Fundus imaging did not show any significant difference between WT and D_2_R KO mice at 3 and 12 months of age (Fig. 6A). Total retinal thickness was not different between WT and D_2_R KO mice at 3 months (161.40± 2.40 μm *vs*. 168.44.02± 14.12 μm, n=4-6; *t*-test, *p* > 0.05, Fig. 6 B, C, D) or 12 months of age (163.14.11± 3.75 μm *vs*. 170.74± 3.10 μm, n=4-6; t-test, *p* > 0.05; Figure 6 B, E, F). At 12 months, both WT and D_2_R KO mice displayed no reduction in the thickness of the different retinal layers in comparison to their younger counterparts (*t*-test, *p* > 0.05 in all cases). Consistent with these results, we did not observe any significant difference between the two genotypes at both ages in the retinal functioning using scotopic and photopic ERG measurements (Fig. 7 A-D, n=4-6; two-way ANOVA, *p* > 0.05).

**Fig 6.**
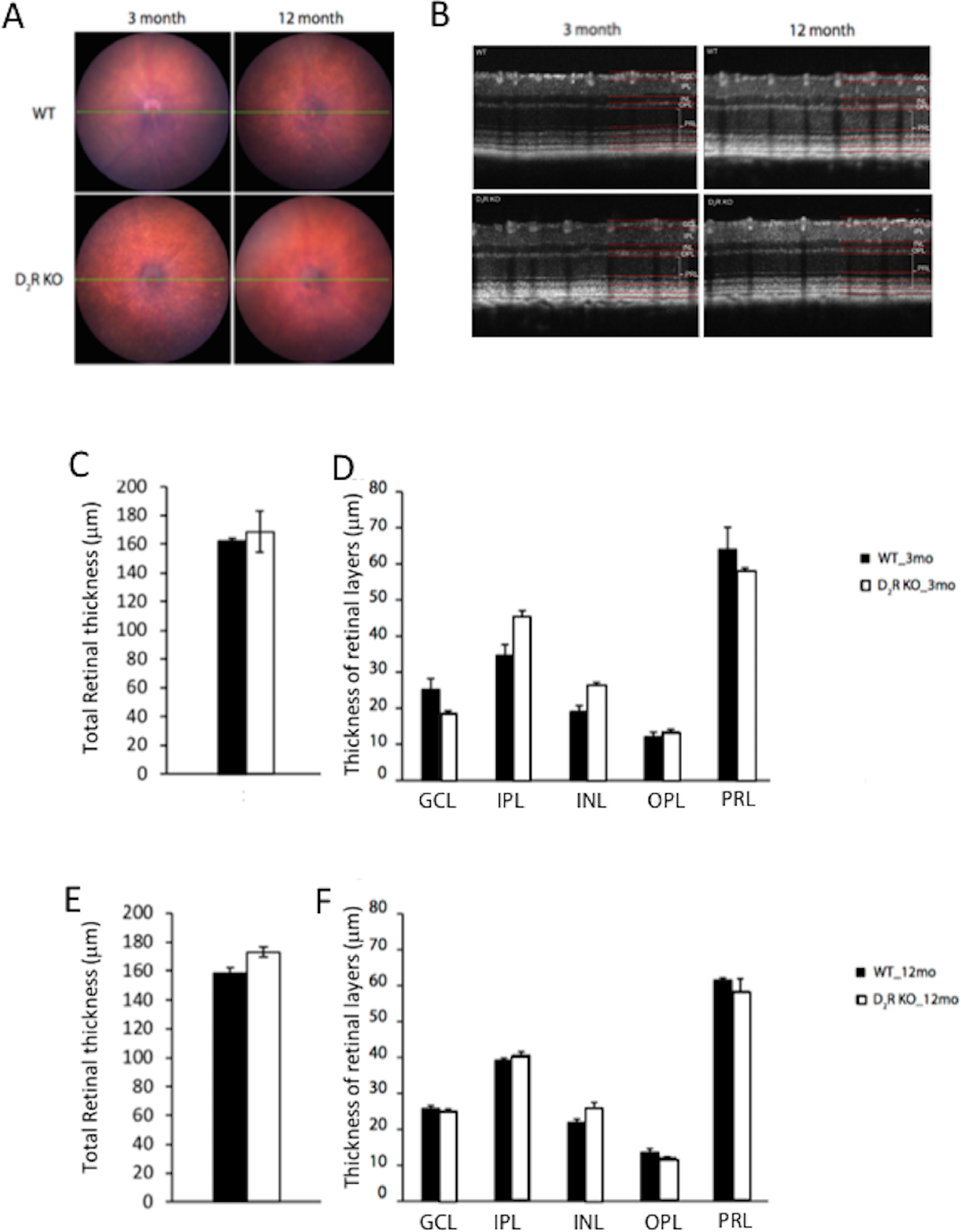
Retinal thickness is unaltered by the lack of D_2_R signaling. (A) Fundus images acquired from WT and D_2_R KO mice of 3 and 12 months of age did not show any significant difference between the two genotypes at both ages. (B) Total retinal thickness was not different between the two genotypes at both the age groups. (C, E). No differences were detected in the thickness of the different retinal layers between the two genotypes at both ages (D, F). GCL: ganglion cells layer, IPL: inner plexiform layer, INL: inner nuclear layer, OPL: outer plexiform, PRL: photoreceptor layer). (*t*-tests, *p* > 0.05 in all cases). Bars represent the means ± SEM. n=4-6.

**Fig 7.**
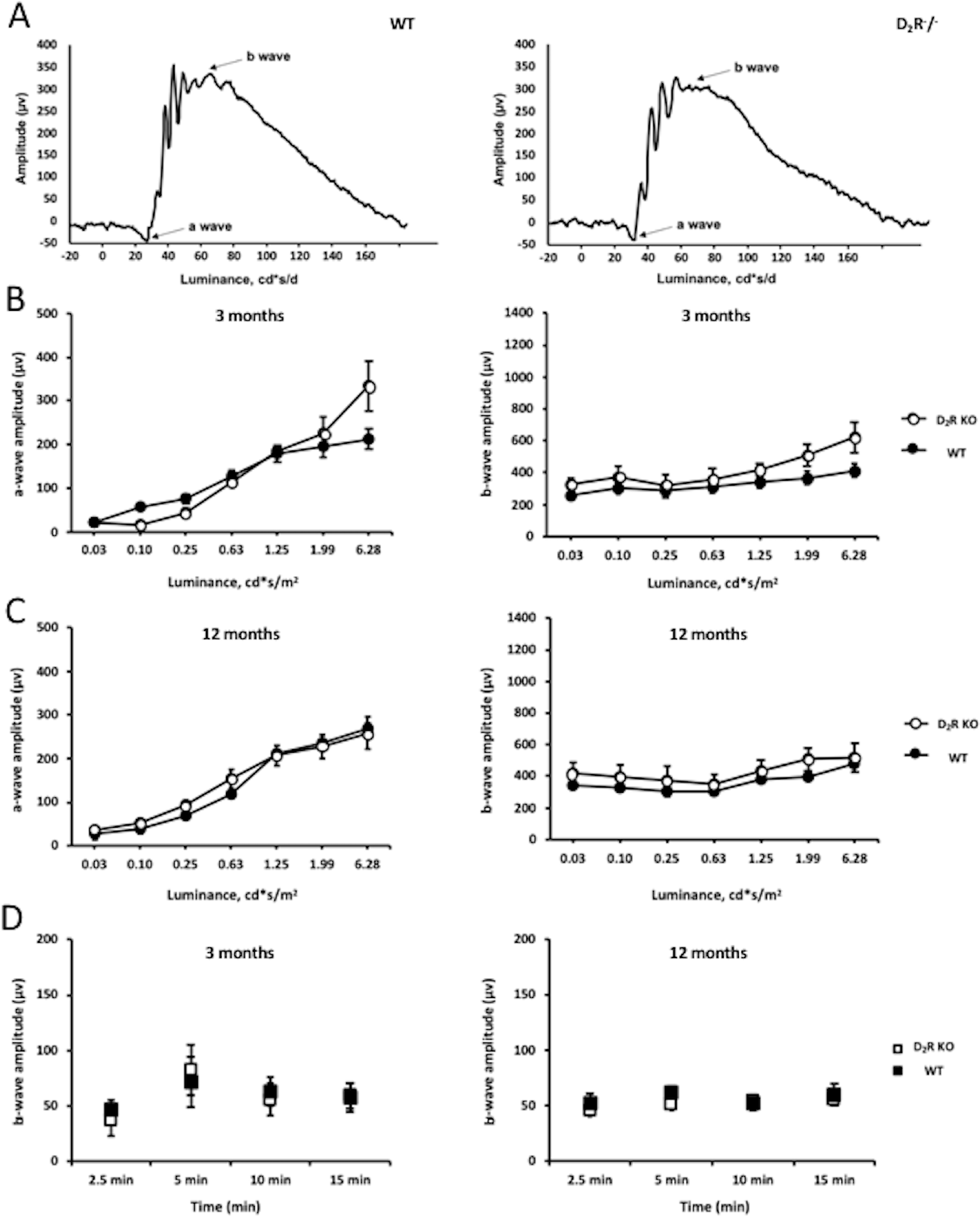
Lack of D_2_R signaling does not affect the scotopic or photopic ERG responses. (A, B) The shape of the waveform in the scotopic ERG did not differ significantly between the WT and D_2_R KO mice. (B, C) The a- and b-wave amplitudes of the scotopic ERGs did not differ between the genotypes both at 3 and 12 months of age. (D) No difference between in the photopic ERG were observed between the WT and D_2_R KO mice at both ages (two-way ANOVA, *p* > 0.1). Circles and squares represent the means ± SEM. n=4-6.

### D_2_R signaling removal has no effects on RPE cell morphology

Morphological analysis via CellProfiler (v2.2.0) of the RPE using ZO-1 immunohistochemistry (Fig. 8A) revealed no difference between the WT and D_2_R KO mice RPE at 3 months of age in parameters like area, compactness, eccentricity and solidity (n=5-6; *t*-test, *p* > 0.05 in all cases, Fig.7 B-I). Further, in the 12 month-old group, we did not find any difference in the parameters mentioned above except for the area of RPE cells which was increased in WT mice compared to D_2_R KO mice (Fig. 8F, *t*-test, *p* <0.05). In WT mice the area was slightly but significantly increased in 12 months old with respect to 3 months old mice (*t*-test, *p* < 0.05), no other differences were observed between young and old WT mice (*t*-test, P. 0.05 in all cases). No differences in the parameters of the RPE morphology were observed between young and old in D_2_R KO mice (*t*-test, p > 0.05 in all cases).

**Fig 8.**
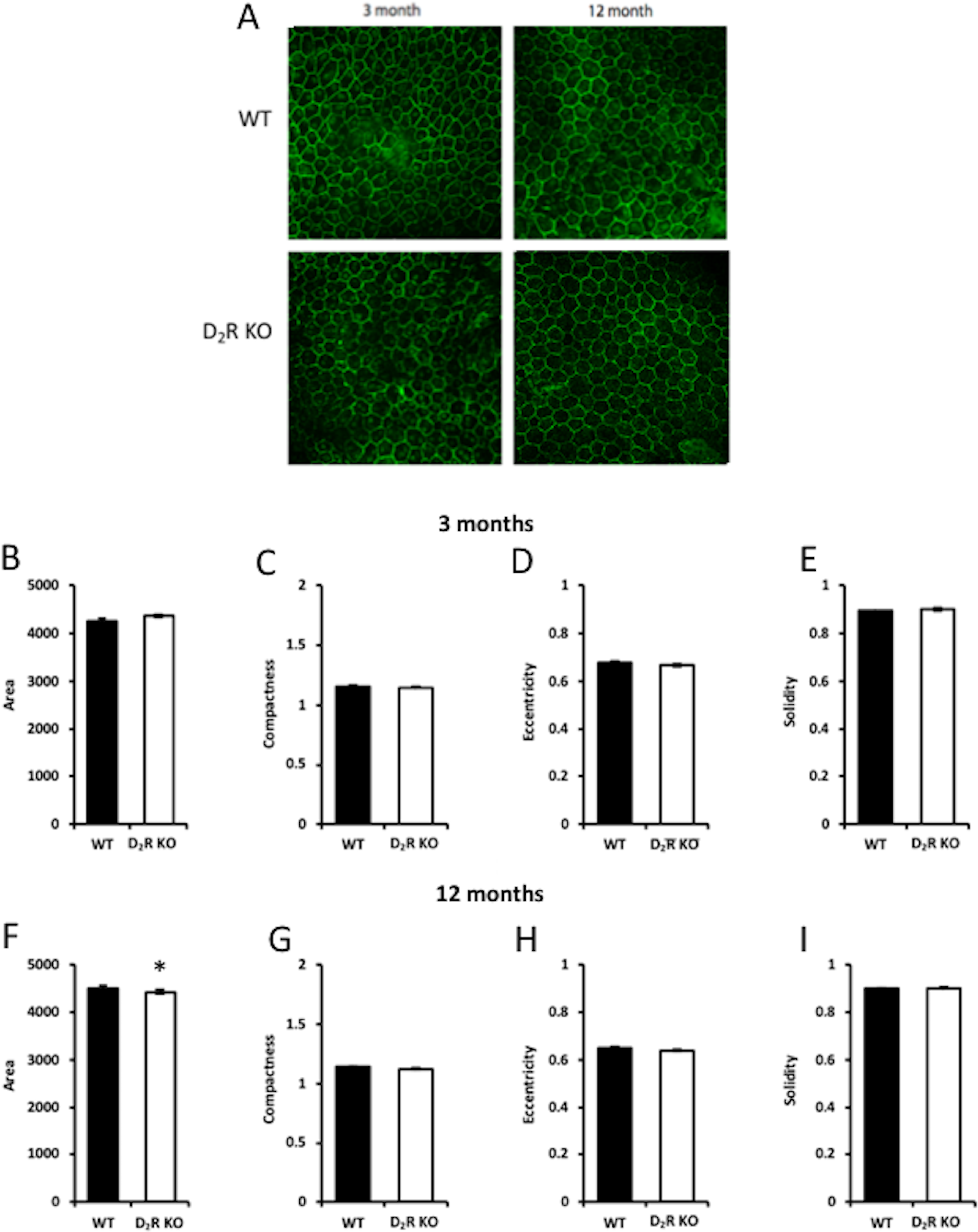
D_2_R signaling does not affect RPE cell morphology. (A) Representative images of ZO-1 staining in WT and D_2_R KO at 3 and 12 months. Analysis with CellProfiler (v2.2.0) indicated that removal of D_2_R signaling did not an affect the morphology of the RPE cells at 3 months of age (B-H) whereas at 12 months of age we observed a slight but significant difference in the total area of RPE cells between WT and D_2_R KO mice (C; *p* < 0.05). Bars represent the means ± SEM. n=4-6.

## Discussion

The burst of phagocytic activity by the RPE after the onset of light has been described in several mammalian species (11,12,13,14,25) and thus many authors have proposed that presence and the timing of the peak are vital for the correct processing of the POS (26, 27). Indeed, experimental data support this hypothesis since β5^−/−^ mice, which lack the peak of the RPE phagocytic activity after the light onset, accumulate lipofuscin and have a reduced visual response during aging (4) even if the total daily phagocytic activity is not different between β5^−/−^ and control mice (4). An additional study has also reported that a small change in the timing of the peak (a phase-advance of about 3 hrs) leads to lipofuscin accumulation (15).

The data reported in this study show: *i*) D_2_R-deficiency completely abolished the burst in phagocytosis activity after the onset of light; *ii*) removal of D_2_R signaling did not affect the expression of canonical clock genes in the RPE; *iii*) analysis of the RPE transcriptome at ZT23 and ZT1 indicated a downregulation of the integrin signaling pathway in D_2_R KO mice; and finally *iv*) D_2_R KO mice did not exhibit any sign of age-related morphological or functional abnormality in the retina. Thus, our data provide evidence that the absence in the burst of phagocytic activity in the early morning does not produce any obvious deleterious effect on the retina or RPE up to one year of age.

The fact that DA signaling plays such an essential role in the regulation of the burst of phagocytic activity is not surprising since DA is a key regulator of daily and circadian function within the retina (28) and it is well known that DA synthesis and release is stimulated by light (29, 30). Similarly, it should not come as a surprise that D_2_R - which are negatively coupled with adenyl cyclase and thus result in a decreased cAMP intracellular level - are involved in the activation of the peak of phagocytic activity by RPE cells since increases in intracellular cAMP decreases phagocytic activity whereas a decrease in cAMP is believed to stimulate phagocytic activity (31). It is also important to mention that dopamine D_4_ receptors do not mediate phagocytosis in the RPE since D_4_R receptor KO mice still show a normal phagocytic activity after the onset of light (32) and activation of these receptors does not shift the RPE circadian clock (20). D_1_R and D_5_R mRNAs are also present in the RPE (20), but the role that these receptors play in the RPE biology is not yet known and thus additional studies are needed to fully understand the role that DA and its associated receptors play in the regulation of RPE biology.

Our data also agree with previous studies (2,4) in which it was reported that activation of FAK signaling is required for the burst in phagocytic activity after the onset of light. We found that the increase in the peak of p-FAK phosphorylation observed in the RPE of WT mice after the onset of light is not present in the RPE of D_2_R KO mouse.

As we have previously mentioned, several studies have shown that the rhythm in phagocytic
activity is under control of the circadian clock (11,12) that is present in the RPE (33, 19, 20, 34). Therefore, it is quite surprising that we did not observe any effect on the daily pattern of clock genes expression in RPE in D_2_R KO mice, thus suggesting that the RPE circadian clock may not be the primary driving mechanism in the signaling cascade controlling the burst in phagocytic activity.

Our RNA-seq data indicate that in both genotypes, a significant number of genes are differentially expressed before and after the onset of light and most of these genes (247) show a significant upregulation in both genotypes. Genes involved in metabolic and biological regulation were most affected and this is not unexpected since the transition from dark to light significantly affect cellular processes.

Using PANTHER, we also discovered that removal of the D_2_R signaling has a significant effect on the genes involved in the integrin pathway thus providing a possible mechanism by which D_2_R signaling and integrin signaling interact to control the daily peak in phagocytic activity. In particular, the observation that *Itgb5* is dramatically down-regulated in D_2_R KO mice suggests that the mechanism that prevents the burst in the phagocytic activity after the onset of light is similar in the D_2_R and β5 KO mice.

Our RNA-seq data also indicate an up-regulation *Pi3K* signaling in WT, as reported by Mustafi et al. (8). However, it essential to note that there are significant differences between our study and the one by Mustafi et al. (8) since in their study they compared the RPE transcriptome between 1.5 and 9.0 hrs after the light onset while we have compared the transcriptome 1 hr before and 1 hr after light onset and thus a direct comparison between our data set and Mustafi’s data set is not possible.

It is worthwhile mentioning that our analysis of the pathways affected by lack of D_2_R is only based on the RNA-seq data and we did not investigate the changes at the proteomic level. Our results indicated a large number of transcripts (see Table 1) are affected by removal of D_2_R and thus it is difficult to select which proteins/pathways should be first investigated. In addition, since the lack of D_2_R signaling does not produce any obvious negative phenotype in the RPE, it is difficult to further investigate D_2_R signaling in the RPE without a clear hypothesis to be tested.

The observation that removal of the daily burst of the phagocytic activity did not produce any major adverse effects on the retina and RPE is somewhat surprising but not completely unexpected since mice lacking MGF-E8 (the ligand for αvβ5 integrin receptor) do not have a peak of phagocytic activity 1-2 hrs after the onset of light (2) and do not show any decrease in scotopic ERG responses at least until 12 months old of age (**Error! Reference source not found.**). While the Finnemann laboratory did not report on the RPE morphology, our data on the RPE morphology suggest that loss of D_2_R signaling and the morning peak in the phagocytic activity do not produce a significant effect in young and old D_2_R KO mice.

A recent study in which phagosome count was perfomerd using electronmicroscopy to image the phagosomes has reported the presence in the mouse of an evening second peak at ZT 13.5 (i.e., h after the onset of dark) (36). In our study we did not extend our analysis at that time of the day since previous studies in C57BL/6, C3H-f^+/+^ and 129/Sv mice - using a similar techique to what we used in our study to count the phagosome - did not report the presence of such a peak (4, 13) and thus we thought that it was unlikely to observe a difference at ZT13.5 between the two genotyoes. In addition, since we did not observe any difference retinal functioning or RPE morphology between the D_2_R and WT genotypes, we also suggest that the presence or absence of the evening peak – as of the morning peak – may not have any significant effect on the retina and RPE.

Altogether these data suggest that the decrease in visual function and the increase in lipofuscin accumulation observed in β5^−/−^ mice is not due to the lack of the phagocytic peak, but rather to other signaling pathways that αvβ5 integrin receptor may modulate. Further studies will be needed to address this critical question and to identify such pathways.

In conclusion, our data demonstrate that DA signaling - via D_2_R - is responsible for the increase in RPE phagocytic activity after the onset of light. D_2_R removal down-regulates integrin signaling and thus FAK phosphorylation at ZT1. Our data also indicate that the absence of the burst of phagocytic in the early morning does not produce any deleterious effects on the retina, at least until 12 months of age.

## Acknowledgments

This work was supported by National Institutes of Health Grants: GM116760 to K.B.; R01EY026291to G.T. and 5U54NS083932 to Morehouse School of Medicine; R01EY004864, R01EY027711, and P30EY006360 to P.M.I.; and an unrestricted departmental grant from Research to Prevent Blindness (Emory Department of Ophthalmology).

